# Decoys and dilution: the impact of incompetent hosts on prevalence of Chagas disease

**DOI:** 10.1101/597708

**Authors:** Mondal Hasan Zahid, Christopher M. Kribs

## Abstract

Biodiversity is commonly believed to reduce risk of vector-borne zoonoses. However, researchers already showed that the effect of biodiversity on disease transmission is not that straightforward. This study focuses on the effect of biodiversity, specifically on the effect of the decoy process (additional hosts distracting vectors from their focal host), on reducing infections of vector-borne diseases in humans. Here, we consider the specific case of Chagas disease and use mathematical population models to observe the impact on human infection of the proximity of chickens, which are incompetent hosts for the parasite but serve as a preferred food source for vectors. We consider three cases as the distance between the two host populations varies: short (when farmers bring chickens inside the home to protect them from predators), intermediate (close enough for vectors with one host to detect the presence of the other host type), and far (separate enclosed buildings such as a home and hen-house). Our analysis shows that the presence of chickens reduces parasite prevalence in humans only at an intermediate distance under the condition that the vector birth rate from feeding on chickens is sufficiently low.

## 1 Introduction

Biodiversity is commonly considered a means for reduction of vector-borne zoonoses risk though it is not always true [1], [2]. Species diversity consists of two elements-species richness: number of species, and species evenness: proportional representation by each species. Adding any host to a vector-host system can reduce or can increase the disease risk. The reduction in disease risk due to the diversity in species is known as the dilution effect. The strength of dilution effect in a system depends not simply on the measures of species richness [3], it also depends on the abundance of dilution hosts relative to focal hosts [4]. The opposite effect is known as the rescue effect when the disease risk is increased. The determination of type of effect is governed by a couple of factors where the competency of the added host is one of the most important ones.

Based on the competency of additional host(s), the effect of distraction of vectors from their suitable host(s) can be broadly divided into two cases – *decoy effect* and *alternative or incompetent hosts’ effect*. Decoy effect involves adding any incompetent (incapable of transmitting the disease) host whereas alternative hosts are capable of transmitting pathogens, but not as much as the focal host. The use of non-human decoys (e.g. livestock) to divert feeding mosquitoes away from humans may reduce vector-borne infections in the short term, but the increase in successful blood meals has the potential to cause long-term increases in mosquito populations and thereby increase the risk of subsequent human exposure [1], [2].

In the last decade, many studies have investigated how biodiversity can help to reduce the incidence of infections of vector-borne zoonoses. Results from many of those studies indicate that it is more difficult than previously thought to predict the effect of biodiversity loss on the spread of vector-borne disease. In 2010, Johnson and Thieltges showed that the strength of dilution effects depends on the relative abundance of dilution hosts relative to focal hosts [4]. Two years later, in 2012, Ostfeld and Keesing suggested that increases in species richness will not always decrease disease risk; indeed, in some cases diversity will cause an increase in infection risk [5]. In 2014, Miller and Huppert (2014) tried to see the effect of host diversity on the prevalence of disease infections [6]. Their study showed the basic reproduction number, *R*_0_, is not necessarily monotonic as a function of species diversity. Thus, the richness in host population can amplify or can dilute disease prevalence depending on vectors’ preference of host. These works challenge the universally established idea that biodiversity always helps to reduce the disease risk. The challenge lies in identifying when and for what types of host–parasite interactions we are likely to find evidence of a negative relationship between diversity and disease.

This study shifts the context from sylvatic to domestic where we study the case of Chagas disease, also known as American trypanosomiasis. This is a potentially life-threatening illness caused by the protozoan parasite, *Trypanosoma cruzi* (*T. cruzi*). It is found mainly in 21 Latin American countries, where it is mostly vector-borne. The vector involved in the transmission of the parasite to humans is a triatomine bug, also known as a ‘kissing bug’. An estimated 8 million people are infected worldwide, mostly in Latin America. It is estimated that over 10,000 people die every year from clinical manifestations of Chagas disease, and more than 25 million people risk acquiring the disease [7]. Cases of Chagas disease have also been noted in the southern United States [8]. According to the World Health Organization (WHO), vector control remains the most useful method to prevent Chagas’ infection in endemic areas[7].

Domestic animals play an important role in the domiciliary transmission of *T. cruzi* [9]. In 1998, Gürtler et al. investigated the influence of humans and domestic animals on household prevalence of *T. cruzi* in vector populations. Their result shows the indoor presence of chickens increases the infected vector density per house [10]. The study did not address directly the impact of presence of chickens on the prevalence of human infections. In 2007, Gürtler et al. studied the role of domestic cats and dogs in *T. cruzi* infection [11]. This study performed an entomological and sero-parasitological survey in two rural villages in Argentina. Both cats and dogs are found as epidemiologically important sources of infection for bugs and householders where dogs are nearly three times more than cats. Gürtler et al. suggested in 1998 that the preventive management of domestic animals is an essential approach to the control of Chagas disease [10]. This suggestion was implemented in 2014 where a community-based intervention was developed based on domestic animal management by De Urioste-Stone et al. and implemented in two cities in Guatemala [12]. This community intervention promoted chicken management as one of the means for reduction of Chagas disease infections.

This study aims to identify conditions, if any, under which the presence of one common domestic animal– chickens–can reduce the vector-human interaction and eventually decrease human disease risk for Chagas. Here chickens are the additional host, which is completely unsuitable for the parasite. So, this work adds to research on species richness, specifically on the presence of an additional host. In this study, we investigate whether this inclusion of an incompetent host (decoy) dilutes or strengthens the force of infection. Chagas disease transmission occurs primarily in rural homes in Latin America. Studies have shown that the practice, common in countries like Argentina, of bringing chickens (brooding hens) into the home for protection of eggs and chicks against predators and then leaving them outside once grown, affects domestic vector populations [13].

Usually, the presence of incompetent hosts reduces the number of encounters between the vectors and the focal host. Eventually it leads us to the perception that this reduces the disease risk. However, some earlier works, where chickens are considered to be in bedroom areas, already proved this perception wrong [9]. The practice among rural areas shows that the residence of chickens changes with time. Thus, the distance between chickens and humans is variable, rather than fixed. This fact motivates us studying the impact of the presence of chickens at varying distances from humans. In our analysis, we consider three different cases depending on the proximity of two hosts, humans and chickens. To analyze these cases, we develop models for transmission separately for each case using dynamical systems.

## 2 Model Development

This work considers three different cases regarding the distance of the incompetent host (chickens) from the focal host (humans): (1) far distance case, (2) intermediate distance case, and (3) short distance case. These cases are determined by the places where chickens are kept by the villagers. Most of the year, villagers keep their chickens either in a place separated from the houses or in some part of their houses. We consider the first of these two scenarios the ‘far distance case’ while we consider the other the ‘intermediate distance case’. However, we consider the scenario ‘short distance case’ when chickens are brought indoors or very close to indoors to ensure their safety at a very young age.

We begin by focusing on mean-field results rather than the range of possible variations, just to see whether the force of infection tends to be strengthened or weakened by the presence of chickens. To do so, we use deterministic models (despite the small populations) since we are interested in qualitative insights. However, since stochastic effects may be significant in small populations, we will also consider a stochastic version of our model(s) to examine possible deviations from the mean.

Vectors’ feeding behavior is very important in modeling vector-borne infectious diseases. In 2006, Ngwa studied the population dynamics of the mosquitos that transmit malaria to humans, incorporating the vector’s feeding behavior into the model. The study divided the vector population in three categories: vectors in the breeding site, vectors moved from the breeding site to human habitat, and vectors moving from human habitat to breeding sites [35]. In a later study, Ngwa et al. further subdivided each of the three categories mentioned above into *N* number of sub-categories assuming that each vector has *N* number of gonotrophic cycles [36]. However, the triatomine vectors of Chagas disease has different behavior than mosquitos in many senses–their reproduction is independent of breeding site, they do not bite in the daytime as mosquitos do, and their movement is very limited compared to mosquitos’. Chagas vectors incline to stay near the sleeping area of the hosts, so vectors’ hiding, or sleeping area is associated with specific host populations. In our model development, we therefore base vector movement and feeding behavior on ideas in research by Gürtler et al. [9, 10].

Most people infected with Chagas disease do not know they have the disease [31, 34]. This happens as the disease is mostly asymptomatic. However, 20% -30% of infected people may develop symptoms at a later stage (chronic stage), but it is too late to cure [30, 32, 33]. Also, people infected with Chagas disease have very limited (less than 1%) access to diagnosis and treatment [29]. Therefore, there is almost no recovery from the disease. Hence, here we consider a SI model in our work. People with Chagas disease can continue their lives without having any symptoms for 10 years or more [29]. So, we assume relatively low disease-induced death rate which allowed us to maintain a constant human population. In order to focus on the effects of the presence of incompetent hosts, we model only two host populations: primary and incompetent. The presence of other competent domestic hosts such as dogs can be incorporated by converting to a transmission-equivalent number of humans using the vectors’ known feeding preferences.

To begin with, we consider the case when chickens sleep in nests separated from the house, either a free-standing hen-house or part of barn or other building (case of far distance). Therefore, whenever bugs start to leave humans for inadequate availability of meals, they can easily and quickly find chickens as a source of their meals. However, here the vectors are unable to anticipate the presence of chickens while they are with humans.

A general compartmental model is used for describing the above mentioned idea mathematically. Here, the two hosts are humans (*H*_1_) and chickens (*H*_2_). Usually, some vectors are associated with humans and others are associated with chickens. However, no infections occur for the vectors (*S*_*v*2_) who bite chickens since chickens are incompetent hosts. The per capita migration rates are independent of hosts’ population density as vectors can not anticipate the presence of hosts due to the distance. Vertical transmission of T. cruzi in humans is already well documented [21, 38], and so we consider this path of transmission in our model, and assume the probability of vertical transmission (i.e., the proportion of offspring of infected mothers which are born infected due to transplacental transmission) for *H*_1_ is *p* and all host demographics are at equilibrium. This is a special case (setting all the parameters related to strain I as zero) of the host switching model of [14]. All these ideas are depicted in Figure 2 and described by the system (1).

**Figure 1:**
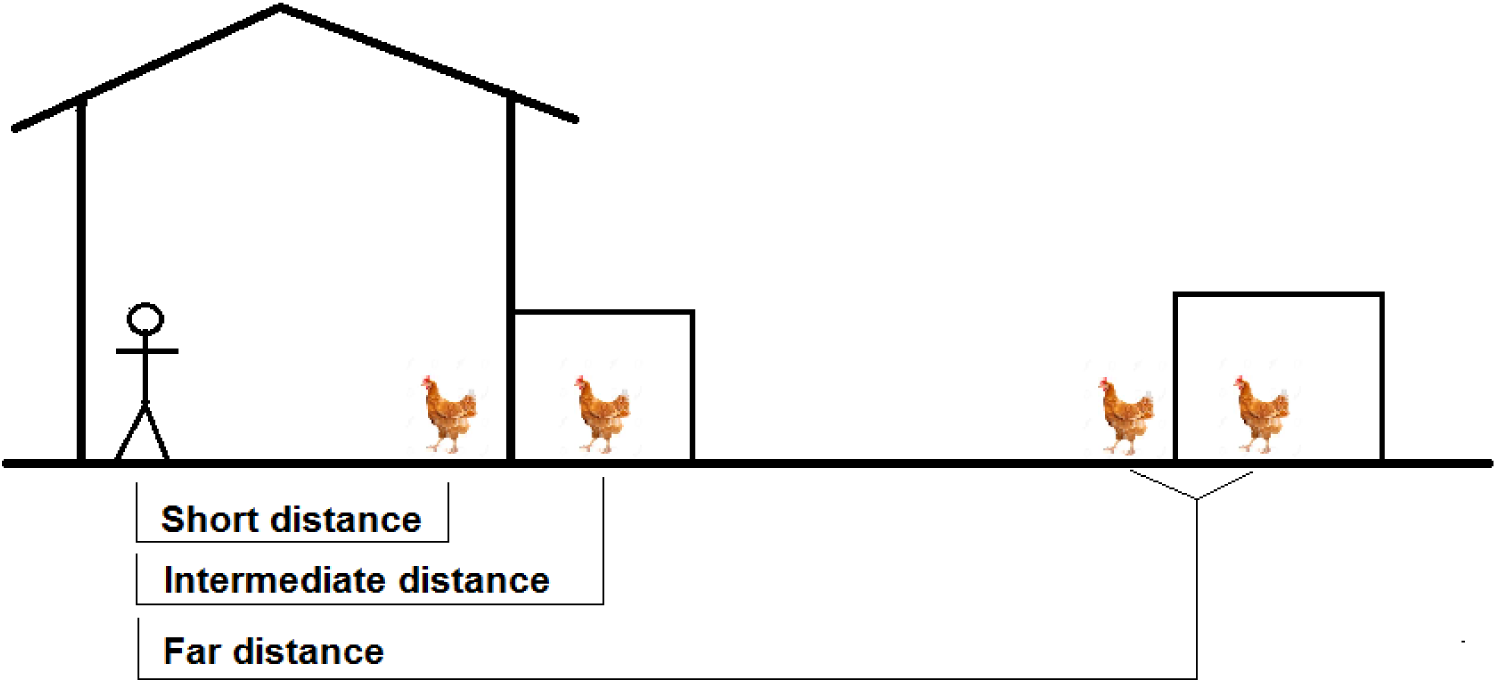
Portrayal of all the three cases.

**Figure 2:**
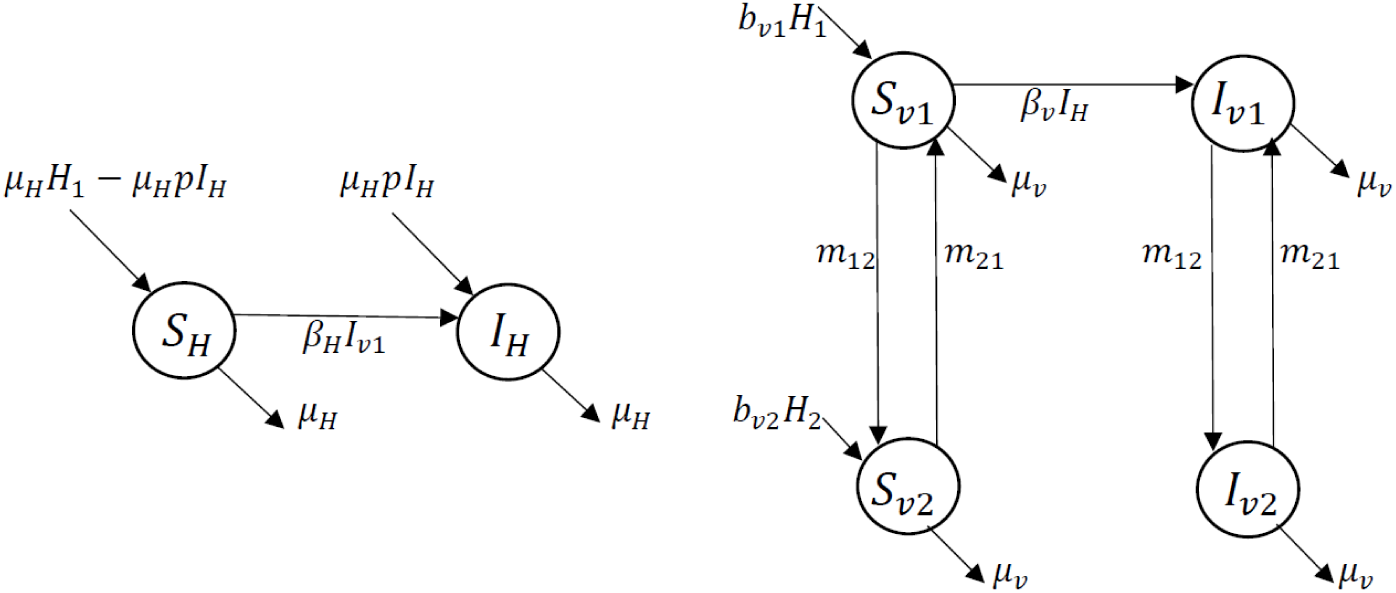
Flow diagram for ‘far distance’, system (1), where movements of vectors are independent of hosts’ density.

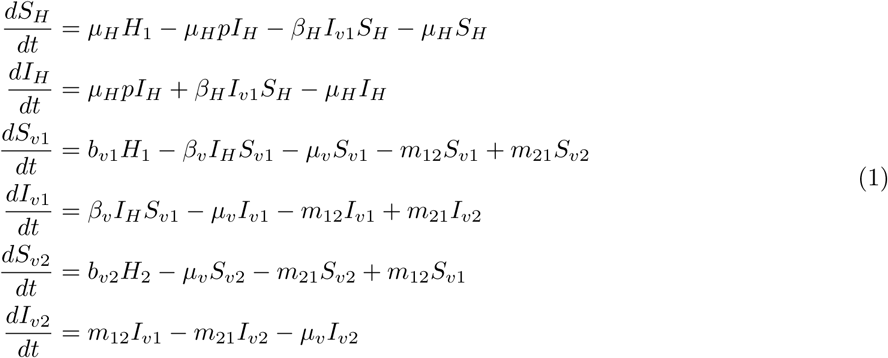

We next consider the scenario when chickens are kept a little bit closer to houses (case of intermediate distance). In this case, chickens live in a hen-house connected to the house, or in a different part of the house than the humans. Here, the proximity allows bugs staying with one host to sense the presence of other hosts and so vectors switch between hosts (humans and chickens) whenever they need. Certainly, the migration rates for vectors between hosts are determined by the availability of blood-meal sources. So, this migration between hosts is dependent on the target host’s density. The model in this case is similar to the previous one, except the migration rates. The per capita migration rates are *m*_12_*H*_2_ for humans to chickens and *m*_21_*H*_1_ from chickens to humans. This case is visualized in Figure 3 and described by the system (2).

**Figure 3:**
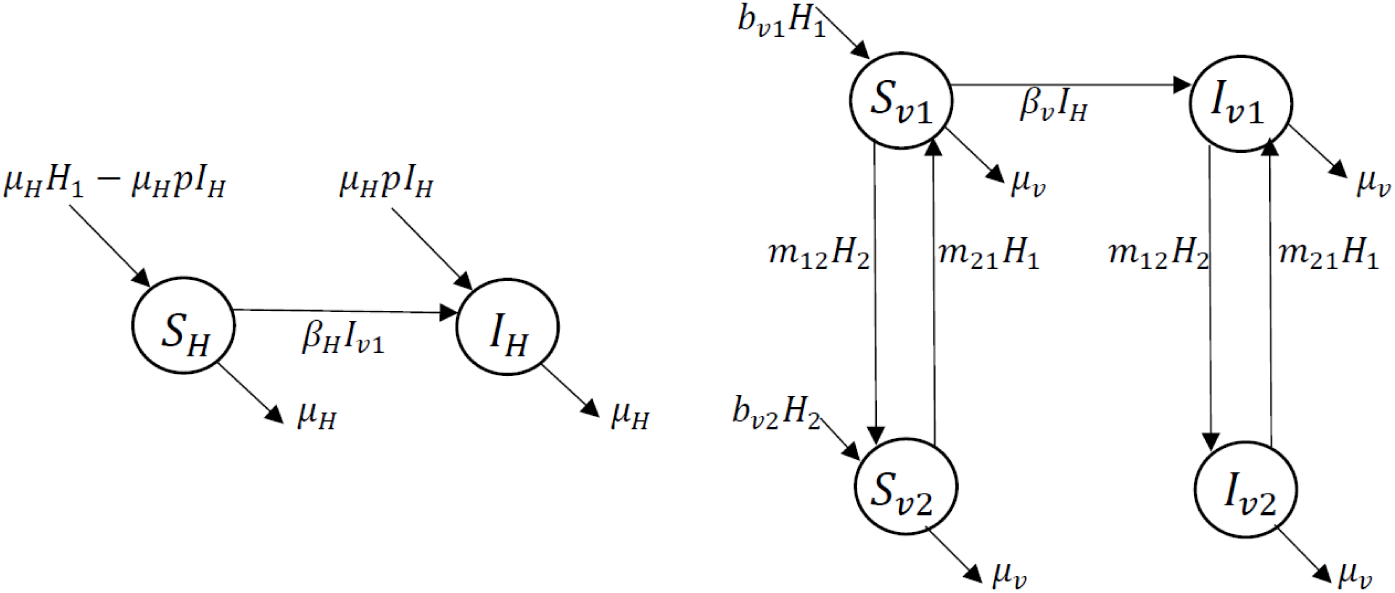
Flow diagram for ‘intermediate distance’, system (2), where movements of vectors are host density dependent.

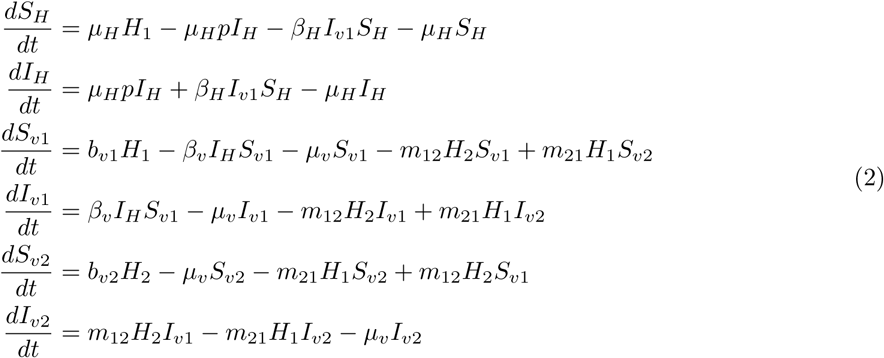

In the last case, chickens are brought so close to humans that vectors do not need to migrate to collect their meals (case of short distance). Now, vectors can bite and take blood meals from whomsoever they want. It is not anymore a host switching case, rather host sharing. So, all the vectors are sharing both of the host populations. Here, we assume that vectors bite humans a proportion *q* of the time. This case is a special case of host sharing model of [14] where all the parameters related to strain I set as zero. This model is portrayed in Figure 4 and represented by the system (3).

**Figure 4:**
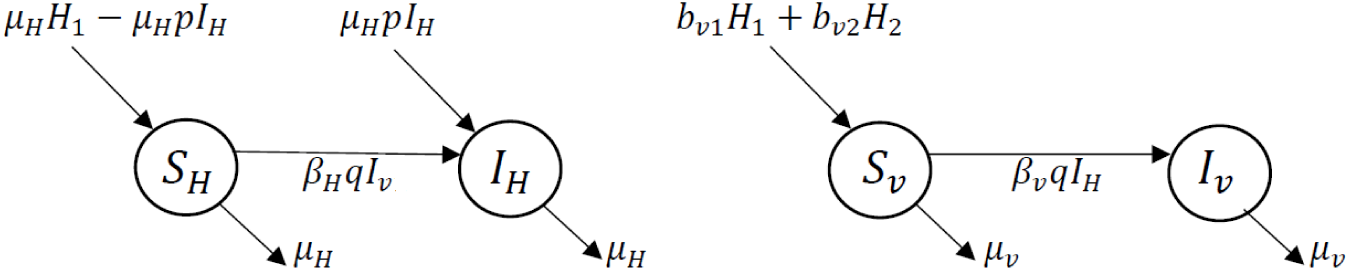
Flow diagram for ‘short distance’, system (3), where vectors don’t need to migrate.

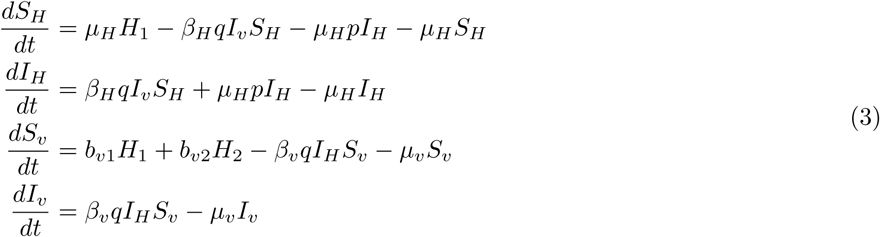

To study the likely variation from mean-field results, we use continuous-time Markov chains (CTMC) as stochastic version of our deterministic model(s). A CTMC model has discrete populations, and discrete events occurring in continuous time as a Poisson process, with expected rates given by the deterministic rates. We use the Gillespie algorithm [37], also known as stochastic simulation algorithm (SSA), to simulate our CTMC model(s).

Table 1 summarizes the variables for all of our models.

**Table 1:** Model variables with definition.

| Variable | Definition |
| --- | --- |
| $S_H$ | Susceptible humans(focal host) |
| $I_H$ | Infected humans |
| $S_{v1}$ | Susceptible vectors associated with humans |
| $I_{v1}$ | Infected vectors associated with humans |
| $S_{v2}$ | Susceptible vectors associated with chickens |
| $I_{v2}$ | Infected vectors associated with chickens |

## 3 Parameter estimation

While estimating parameters, we tried to take the values from the same geographical context (Argentina) to make our analysis more appropriate. Some of these parameter estimates are very rough, and we include them here primarily in order to generate illustrative qualitative trends. This study considers *Triatoma infestans* as the vector since this is the most common vector of *T. cruzi* in South America, including Argentina [15], [16], [17].

During our careful literature review, we did not find any documented data for infection rates for humans and for vectors (*β*_*H*_ and *β*_*v*_ respectively). To estimate these values we used the method from [18] which gives the following formulas for our case:

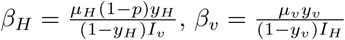

where *y*_*H*_ and *y*_*v*_ represent the prevalence of the disease in humans and chickens respectively. We take 27.81% (*y*_*H*_) for humans [19] and 4.1% (*y*_*v*_) for vectors [20], and multiply the household size and the number of bugs in a house by these prevalence values to find the value of *I*_*H*_ and *I*_*v*_. In our literature review, we found the value 0.09 (documented as 9%) [21] for probability (proportion) of vertical transmission (*p*). For the human death rate, (*µ*_*H*_) we take the reciprocal of their average lifespan and 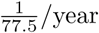 [22]. However, we did not find any direct documented data for vectors’ death rate (*µ*_*v*_). So, we used different data from the study done in 2015 by Medone et al. [15] and did our own estimation to find average lifespan for *Triatoma infestans* [Table 2] and finally take the reciprocal to get 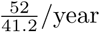 as value for *µ*_*v*_. Finally, using our own formula the infection rates are obtained as

**Table 2:** Estimation of average lifespan for *Triatoma infestans* while feeding only on humans and chickens (base data are taken from [15])

| Feeding pattern | Stage | Duration | Lifespan by gender | Lifespan by host's | Mean lifespan |
| --- | --- | --- | --- | --- | --- |
| Fed on humans | Egg to Nymph V | 29.2 wks | 45.5 wks | 46.05 wks | 41.2 wks<br>($\frac{41.2}{52}$ year) |
|  | Adult as male | 16.3 wks |  |  |  |
|  | Egg to Nymph V | 29.2 wks | 46.6 wks |  |  |
|  | Adult as female | 17.4 wks |  |  |  |
| Fed on chickens | Egg to Nymph V | 18.9 wks | 34.0 wks | 36.35 wks |  |
|  | Adult as male | 15.1 wks |  |  |  |
|  | Egg to Nymph V | 18.9 wks | 38.7 wks |  |  |
|  | Adult as female | 19.8 wks |  |  |  |

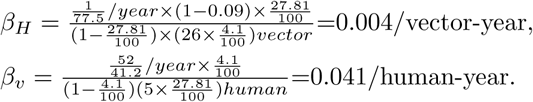

In our literature review, we did not find any documented data for vectors’ birth rate per human (*b*_*v*1_). Hence, we used the total vector population in disease free state from Table 4 to do back-calculation for estimating *b*_*v*1_. Setting migration rates (*m*_12_ and *m*_21_) as zero in 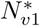 for intermediate case, we get 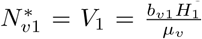 and eventually we get the formula:

**Table 3:** Summary of estimated model parameters.

| Par. | Definition | Value | Units | Reference |
| --- | --- | --- | --- | --- |
| $\beta_H$ | Infection rate for human | 0.004 | 1/vector-year | This study |
| $\beta_v$ | Infection rate for vectors | 0.041 | 1/human-year | This study |
| $p$ | Probability of vertical transmission in humans | 0.09 | - | [21] |
| $b_{v1}$ | Vectors birth rate (per human) | 2.95 | vector/human-year | This study |
| $b_{v2}$ | Vectors birth rate (per chicken) | 14.75 | vector/chicken-year | This study |
| $\mu_H$ | Death rate for human | 1/77.5 | 1/year | [22] |
| $\mu_v$ | Death rate for vectors | 52/41.2 | 1/year | This study |
| $m_{12}$ | migration rate from humans to chickens in (2) | $\frac{365}{(14 \times 15)}$ | 1/chicken-year | This study |
| $m_{21}$ | migration rate from chickens to humans in (2) | $\frac{365}{(14 \times 5)}$ | 1/human-year | This study |
| $q$ | proportion of time at which vectors fed on humans | 1/6 | - | [24] |

**Table 4:** 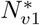 for all three cases.

| Far distance | Intermediate distance | Short distance |
| --- | --- | --- |
| $\frac{b_{v1}H_1 + b_{v2}H_2 \left( \frac{m_{21}}{m_{21} + \mu_v} \right)}{\mu_v \left( 1 + \frac{m_{12}}{m_{21} + \mu_v} \right)}$ | $\frac{b_{v1}H_1 + b_{v2}H_2 \left( \frac{m_{21}H_1}{m_{21}H_1 + \mu_v} \right)}{\mu_v \left( 1 + \frac{m_{12}H_2}{m_{21}H_1 + \mu_v} \right)}$ | $\frac{b_{v1}H_1 + b_{v2}H_2}{\mu_v}$ |

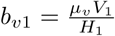

Our study found documented value for household size as 5 persons [23] and for bugs per infested house as 26 (1429 bugs in 55 houses, only the domiciliary cases are considered since we are looking for vectors’ birth rate per human) (*V*_1_) [9]. The vector data were taken from houses where other hosts (dogs and cats) live also. In our literature review, we got 2.0 dogs and 0.5 cats per house [23]. So, to make the value of *b*_*v*1_ truly per human we use the equivalence relation (based on the vectors’ feeding pattern) among hosts done by Gürtler et al. [24] where they show one dog or cat is equivalent to 2.45 (mean of 2.3 and 2.6) humans. After doing some basic arithmetic, we found the equivalent number of persons per household is 11.125 (we use this as *H*_1_ only for the estimation of *b*_*v*1_, otherwise we used 5 as the value of *H*_1_). Using this equivalent value in the above formula for *b*_*v*1_ we obtained

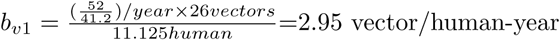

Since vectors fed on chickens five times more than humans [24], we multiply the value of *b*_*v*1_ by 5 to get the value for *b*_*v*2_ which gives 14.75/chicken-year.

For estimating migration rate from chickens to humans (*m*_21_), we take the time duration of vectors’ last feeding to seeking a new host from [25], convert it to year, take the reciprocal of it and finally divide by household size which gives 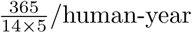. For estimating the value of *m*_12_, we similarly use the number of chickens/household, which is 15 [23] and get 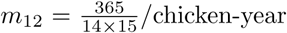. For the proportion of time at which vectors fed on humans (*q*), we found 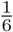 (documented as five times more fed on chickens compare to humans) [24]. All the parameter estimates are summarized in Table 3.

## 4 Analysis

The goal of this study is to analyze the impact of the additional incompetent host on the prevalence of Chagas disease among humans. The equilibria and the basic reproduction number (*R*_0_) are primary indicators for such observations.

To find the equilibria of all three dynamical systems, we set every single equation equal to zero for each model separately and solve. In this process, we find the total vector population 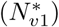 from the disease-free equilibrium; those are shown in Table 4. However, expressions provided in Table 4 hold for both the disease-free and endemic equilibria, because all three models assume that infection does not affect vector birth or death rates. We also get the infected human population 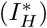 from the endemic equilibrium. Even though we are interested in observing the behavior of the infected population class, we still need to know the basic reproduction number (*R*_0_) as it plays a very important role in interpreting the behavior of any infectious disease. To find the expression for *R*_0_ we use the next generation method [26]. The expressions for *R*_0_ and 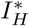 for all three cases are in Table 5.

**Table 5:** *R*_0_ and 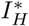 for all three cases, note 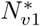 is a function of *H*_2_ in each case.

| Case | $R_0$ | $I_H^*$ |
| --- | --- | --- |
| Far distance | $\frac{p}{2} + \sqrt{\frac{p^2}{4} + \frac{\beta_H \beta_v H_1 N_{v1}^*}{\mu_h \mu_v \left( 1 + \frac{m_{12}}{m_{21} + \mu_v} \right)}}$ | $\frac{-\mu_H (1-p) \mu_v \left( 1 + \frac{m_{12}}{m_{21} + \mu_v} \right) + \beta_H \beta_v H_1 N_{v1}^*}{\beta_v [\mu_H (1-p) + \beta_H N_{v1}^*]}$ |
| Intermediate distance | $\frac{p}{2} + \sqrt{\frac{p^2}{4} + \frac{\beta_H \beta_v H_1 N_{v1}^*}{\mu_H \mu_v \left( 1 + \frac{m_{12}H_2}{m_{21}H_1 + \mu_v} \right)}}$ | $\frac{-\mu_H (1-p) \mu_v \left( 1 + \frac{m_{12}H_2}{m_{21}H_1 + \mu_v} \right) + \beta_H \beta_v H_1 N_{v1}^*}{\beta_v [\mu_H (1-p) + \beta_H N_{v1}^*]}$ |
| Short distance | $\frac{p}{2} + \sqrt{\frac{p^2}{4} + \frac{\beta_H \beta_v H_1 q^2 N_{v1}^*}{\mu_H \mu_v}}$ | $\frac{-\mu_H (1-p) \mu_v + \beta_H \beta_v H_1 q^2 N_{v1}^*}{\beta_v [\mu_H (1-p)q + \beta_H q^2 N_{v1}^*]}$ |

The expressions for *R*_0_ and 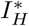 clearly manifest that 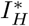 is positive in all three cases iff *R*_0_ >1. Now, to check the impact of the presence of our incompetent host, chickens (*H*_2_), we define 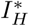 as a function of *H*_2_ and then take the derivative of this newly defined function with respect to *H*_2_. The expressions of these derivatives for far distance and short distance cases are given in Table 6. From the expressions, it is evident that these derivatives are always positive, which implies bringing chickens into the system always makes the situation worse for humans.

**Table 6:**
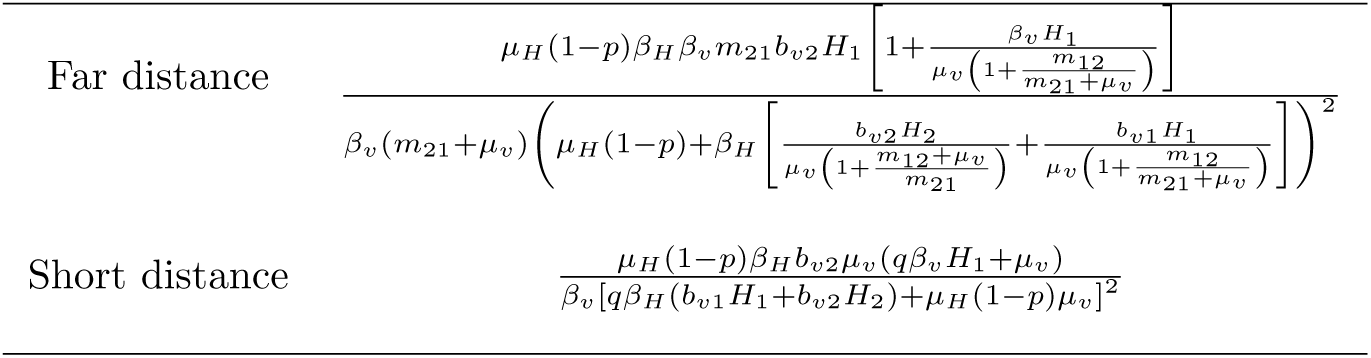
Derivatives of 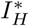 with respect to *H*_2_.

However, the consequences for the intermediate distance case are not straightforward. Here, the value of the derivative 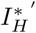 (with respect to *H*_2_) either can be positive or can be negative depending on certain conditions. In our analysis, we find housing chickens at an intermediate distance from humans can cause the prevalence of Chagas disease among humans to be slowed down only if

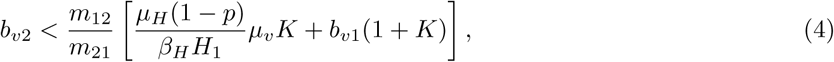

Where 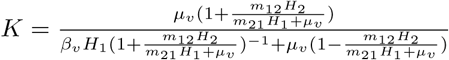.

The above condition (4) on *b*_*v*2_ can only be true if

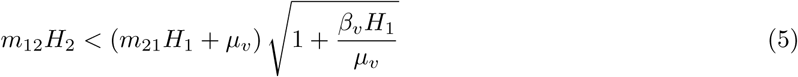

Here, the second addend in (4) is directly proportional to *b*_*v*1_, and the first term is inversely proportional to both *β*_*v*_(through *K*) and *β*_*H*_. Thus this condition is easy to satisfy when vectors have easy access to humans (high *b*_*v*1_) or disease transmission (*β*_*H*_ and *β*_*v*_) is low. So, the presence of chickens is helpful in this case if the birth rate of vectors with chickens is less than a certain threshold value which is relative to the birth rate of vectors with humans and inversely proportional to the infection rate among humans.

A local sensitivity analysis of the potential endemic prevalence of Chagas disease 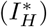 and *R*_0_ was performed (Figure 5). Sensitivity indices for quantities were carried out for all model parameters, and the outcomes indicate that neither quantity is highly sensitive to those model parameters which are more difficult to estimate well. Both measures 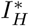 and *R*_0_ are most sensitive to vector longevity, *µ*_*v*_, which is a well known quantity. All normalized sensitivity indices for *R*_0_ except *µ*_*v*_’s were less than 1/2. Remarkably, neither measure (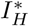 and *R*_0_) is highly sensitive to vector (feeding on humans) birth rate (*b*_*v*1_). These sensitivity analyses show that the parameters not known well are less influential and the most influential parameters are known well. Thus, the results of this study will not be significantly affected even if the actual values of our estimated parameters vary significantly from our estimation.

**Figure 5:**
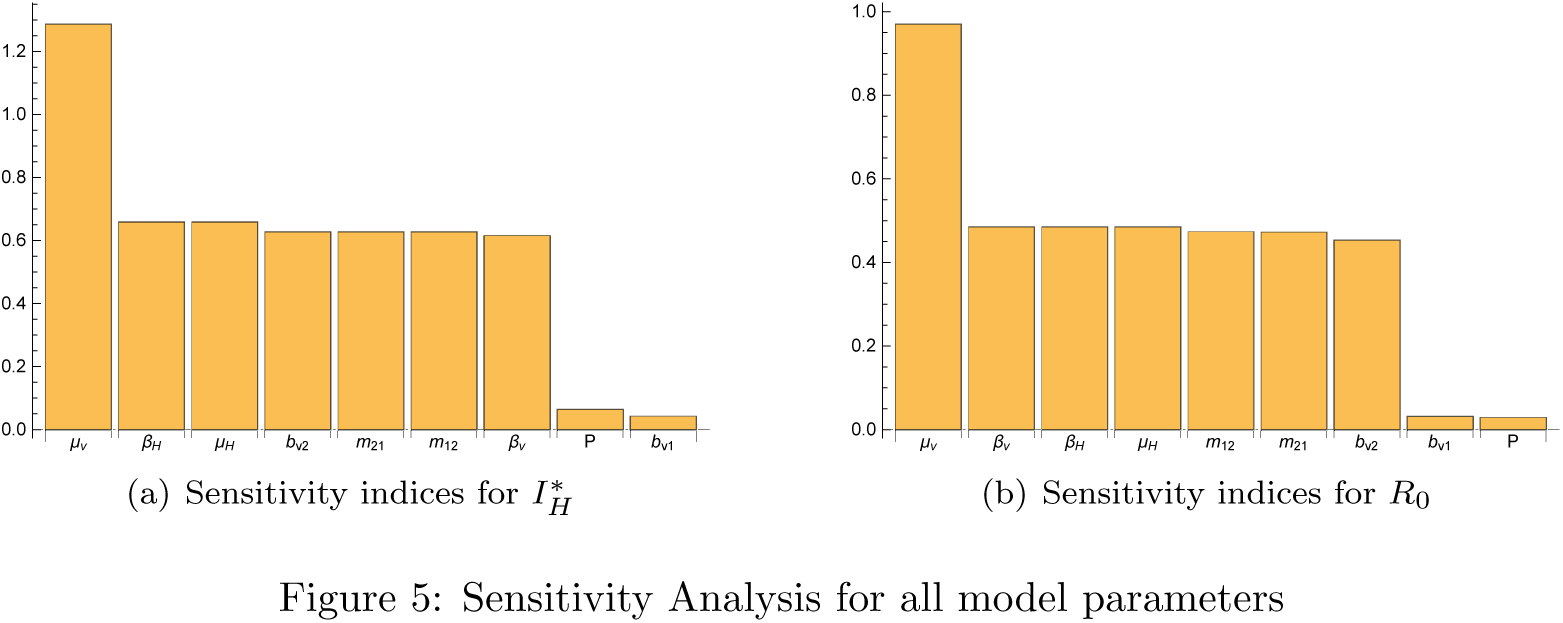
Sensitivity Analysis for all model parameters.

To facilitate interpretation, here we illustrate our results numerically for only the helpful case (intermediate case) in brief. At baseline (our estimated parameter values), we get *R*_0_ = 1.58 and we also find the condition *b*_*v*2_ < 19.57/chickens-year at which the presence of chickens is helpful in reducing prevalence of Chagas disease in humans. In our analysis, we find *R*_0_ strictly decreasing function of *m*_12_ and strictly increasing function of *m*_21_. However, *R*_0_ increases for up to a certain number of chickens and then start to decrease (Figure-6). This implies that for our parameter values the presence of chickens can reduce the infections in humans depending on the number of this incompetent host. Now, the condition for making the presence of chickens helpful becomes easier to satisfy as migration of vectors from humans to chickens increases and it becomes difficult as migration from chickens to humans increases. However, increasing the number of chickens makes the helpfulness criterion (4) easier to satisfy. All the numerical values here are based on our parameter estimations which can be different with other set of parameter values. However, the qualitative result will be the same regardless of parameter values.

We use a deterministic model to develop qualitative insights into our systems. A stochastic model such as a CTMC shows how randomness in individual behavior causes deviations from mean-field results. Here we show results from a CTMC for the intermediate distance case only since the presence of chickens in the other two cases never reduces human infections. To analyze the CTMC model, we performed 100 simulations for each value of *H*_2_. Figure 7 plots the number of infected people at endemic equilibrium 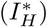, where the solid line indicates the mean-field results for the intermediate case, and the dashed lines represent an envelope of ±1SD. This figure clearly illustrates that the CTMC simulation results follow the same trend established in the deterministic model; thus, variation due to stochasticity does not oppose the answer provided by the deterministic model to the central question of this study.

**Figure 6:**
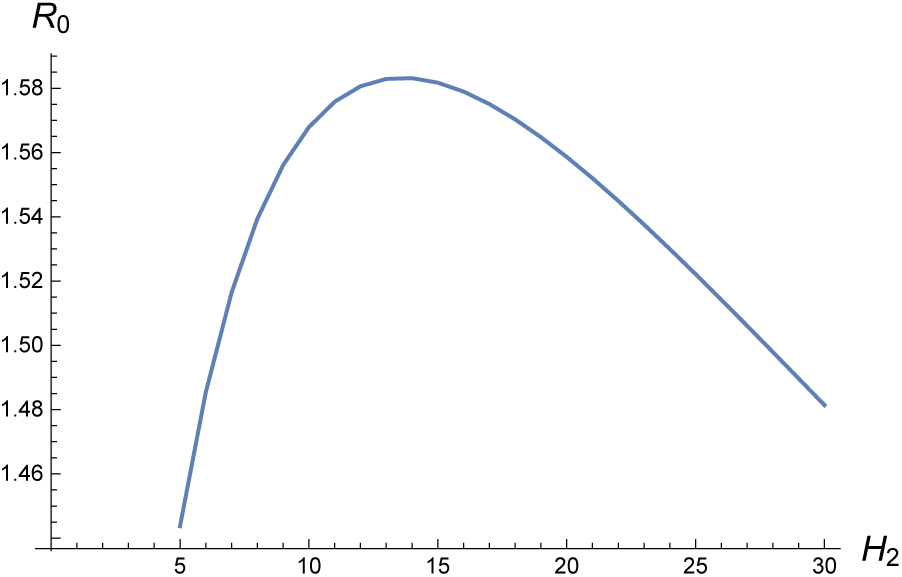
Behavior of *R*_0_ as number of chickens (*H*_2_) varies.

**Figure 7:**
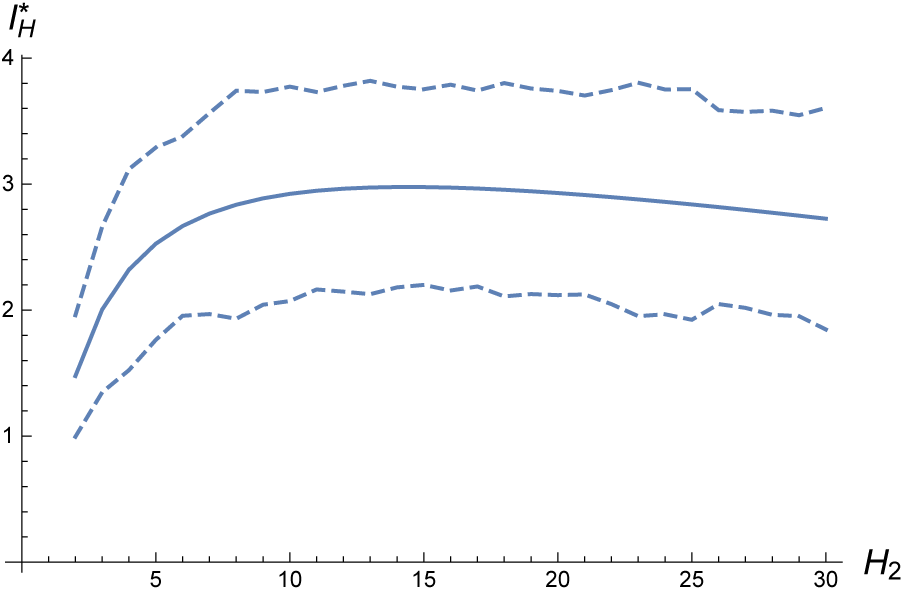
The solid curve shows the mean-field value of the endemic equilibrium; dashed curves indicate an envelope of ±1SD deviation from that value, based on simulations from the CTMC

Our results and analyses show that the presence of an incompetent host, in our case chickens, can reduce the prevalence of Chagas disease in humans under certain conditions only if chickens are placed at an intermediate distance from humans.

## 5 Discussion

This is the first study to understand how the placement of chickens in households affects the transmission of Chagas disease in humans. The case when farmers bring their chickens inside the bedrooms, or very close to bedrooms, increases the number of infections in humans. This short-distance case was studied by Gürtler et al. [10] who focused on vector infections, rather than human infections. Even though the goal of this study is completely different from theirs, we show here analogous computations for comparison. Our analysis shows that both the number, and density of infected vectors increase when chickens kept indoors, after a brief transient decrease (Figure 8). This result agrees with the result of [10] in terms of vector infection. The other two cases, intermediate distance and far distance, studied here are new approaches to understand the impact of chickens’ presence in households.

**Figure 8:**
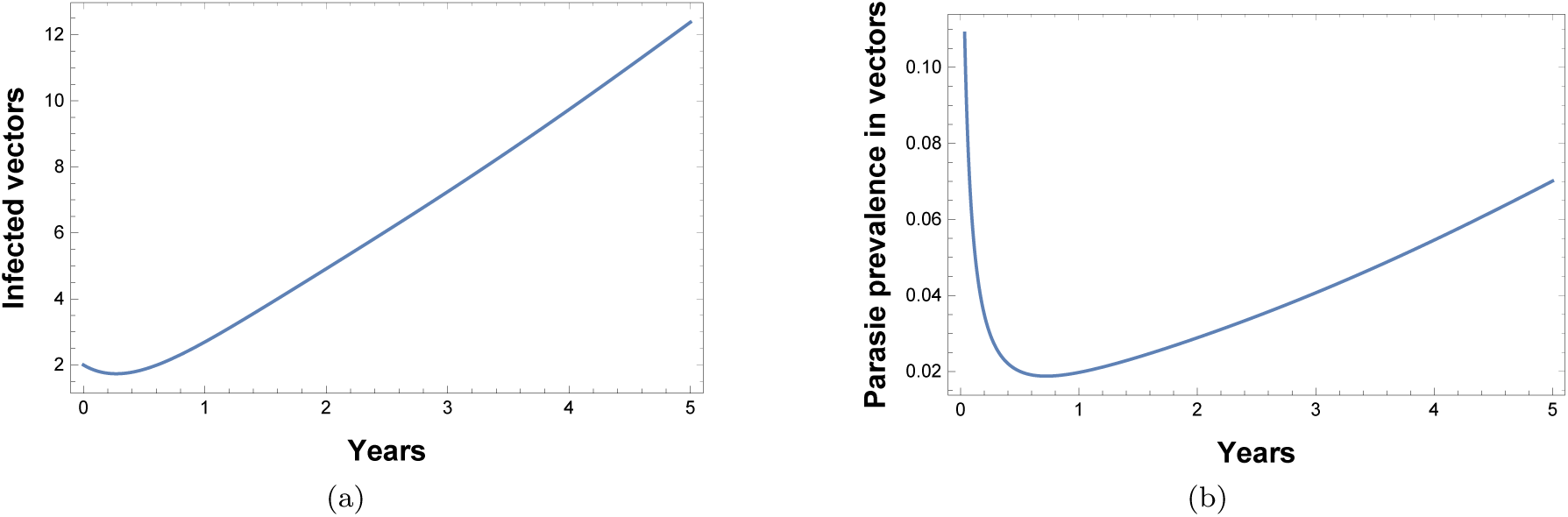
Infections in vectors when chickens are kept indoors (Short distance case)

For the far case, where vectors cannot anticipate the location of chickens, the decoy process does not help to reduce human infections. Here, vectors try to stay with humans as long as they can survive since they can’t see any alternative food sources around them. So, by the time when a portion of them start to leave humans, the infections are already spread among humans at a large scale. Consequently, this case is not helpful for the purpose of controlling the prevalence of infections among humans.

In the remaining case, when chickens reside at a distance (adjacent to humans) such that vectors can detect the presence of remaining host while staying with the other, vector populations begin to migrate from humans to chickens in search of their blood-meals. The vector population with chickens will increase with time for having enough food sources and at some point they will start to move towards humans in search of new blood-meal sources. The net effect of vectors’ migration from humans to chickens, and from chickens to humans will determine the effects of chickens’ presence. Our results show that there are certain conditions under which human infections can be reduced. This will happen as most of the vectors will switch from humans to chickens before people in houses are infected that much.

The presence of chickens in houses can only help to reduce the prevalence of Chagas disease among humans when villagers keep their chickens at a distance which allows the vectors to anticipate the location of other hosts, but does not allow vectors to share both of the chickens and humans as their blood meal sources. Hence, it can be concluded that the decoy process, by the presence of an incompetent host, does not always help to reduce the disease prevalence among humans.

Results here offer valuable information to contribute in improving control of Chagas transmission. However, proper understanding of the outcomes of this study depends on the distance from which vectors can sense the presence of hosts. Triatomine vectors detect host by identifying the presence of factors such as, water vapor, heat, and distinctive odors from different odorants (including *CO*_2_)[27, 28]. We found only one documented data source which says triatomine bugs can identify human presence from two meters by detecting heat [28]. Thus research is needed to identify true threshold distance between humans and chickens to distinguish between short distance and intermediate distance cases. As an NTD, Chagas disease has very few data on its transmission cycles. Consequently, we use relatively simple models based on available information regarding demographics and transmission mechanisms, and parametrized by few data what are available. Reliable estimations of our model parameters will better ground our quantitative results. In addition, availability of additional key rates relating to transmission will permit more detailed models.

## 6 Acknowledgment

We thank Dr. Ricardo E. Gürtler, Professor and Laboratory Head, Department of Ecology, Genetics and Evolution, University of Buenos Aires, Argentina, for providing necessary information regarding our few queries which helped us to prepare part of our manuscript and also for suggesting an appropriate reference to our study.

## Notes

#### Summary of Updates

Introduction: Literature review part revised significantly to connect it to our research question better; Analysis: #1. We also rewrite the paragraph relating sensitivity analysis to make more sense. #2. We add the analysis of an stochastic version (CTMC) of our deterministic model to study the likely variation from mean field results. In this connection, we also added the Figure 7. Discussion: #1: First paragraph in the "Discussion" section is added to compare our results with a similar earlier study (1998). Related graph is provided also. #2: The remaining of this section is rewritten significantly to discuss each of three cases in an individual paragraph. #3. Also, we include some limitations of our study at the end of the paper.

